# Using neural distance to predict reaction time for categorizing the animacy, shape, and abstract properties of objects

**DOI:** 10.1101/496539

**Authors:** J. Brendan Ritchie, Hans Op de Beeck

**Author notes:** Corresponding author: J. Brendan Ritchie Address: Psychologisch Instituut, Tiensestraat 102 – box 3714, 3000 Leuven, Belgium.

## Abstract

A large number of neuroimaging studies have shown that information about object category can be decoded from regions of the ventral visual pathway. One question is how this information might be functionally exploited in the brain. In an attempt to answer this question, some studies have adopted a neural distance-to-bound approach, and shown that distance to a classifier decision boundary through neural activation space can be used to predict reaction times (RT) on animacy categorization tasks. However, these experiments have not controlled for possible visual confounds, such as shape, in their stimulus design. In the present study we sought to determine whether, when animacy and shape properties are orthogonal, neural distance in low- and high-level visual cortex would predict categorization RTs. We also investigated whether a combination of animacy and shape distance might predict RTs when categories crisscrossed the two stimulus dimensions, and so were not linearly separable. In line with previous results, we found that RTs correlated with neural distance, but only for animate stimuli, with similar, though weaker, asymmetric effects for the shape and crisscrossing tasks. Taken together, these results suggest there is potential to expand the neural distance-to-bound approach to other divisions beyond animacy and object category.

## 1. Introduction

Using multivariate pattern analysis (MVPA) several neuroimaging studies have investigated object category-selectivity in regions of the ventral visual pathway ^1^. At the same time, some findings have been interpreted as showing that apparent categorical effects in high-level regions of the ventral pathway may be accounted for by more low-level visual features of stimuli ^2–4^. Only recently have studies begun to employ stimulus designs that tease apart the separate contribution of object category and other visual properties, such as shape, to the neural response in ventral pathway brain regions ^5–7^. A virtue of the stimulus designs of these studies is that, by making category and shape orthogonal properties of object images, they help rule out the possibility that the categorical effects in later regions of the ventral pathway are accounted for by low-level visual features of stimuli ^8^.

Besides possible visual confounds, another issue is that evidence of category-related information in a brain region does not guarantee that it is functionally exploited and therefore represented. A first step in trying to address this issue is to show that decodable information can be used to predict behavior ^9–11^. One way of moving beyond this first step is to attempt to not just predict, but also partially explain, observer behavior using a neural distance-to-bound analysis (NDBA), which involves correlating distance from a classifier decision boundary in activation space with observer categorization reaction times (RTs). Based on the predictions of classic psychophysics ^12,13^, if the neural activation space reflects the representation that determines observe categorization response, then RTs should negatively correlate with distance from the decision boundary ^14^. Recent studies have used NDBA to predict observer RTs for animacy categorization tasks based on neural distances measured with human fMRI and MEG ^15–18^.

To date, none of the studies employing NDBA have used stimulus designs that dissociate object category from other visual features of object images. In the present study we sought to determine whether, when animacy and shape properties are orthogonal, neural distance in low- and high-level visual cortex would still predict observer RTs on an animacy categorization task. We also sought to assess whether neural distance might predict RTs when observers categorize object images based on shape. Finally, we also investigated whether a combination of animacy and shape distance might predict RTs when task categories crisscrossed the two stimulus dimensions, and so were not linearly separable along these dimensions.

## Materials and Methods

### Participants

15 adult volunteers (8 Female; mean age = 30; right-handed) participated in the experiment. All participants had normal or corrected vision, provided written informed consent, and were financially compensated for their participation. The experiment was approved by the Medical Ethics Committee of UZ/KU Leuven and all methods were performed in accordance with the relevant guidelines and regulations.

### Stimuli

Stimuli were 32 natural images of objects, converted to greyscale with scene background removed (Figure 1A). All images were cropped to 700 × 700 pixels and subtended ~ 10° of visual angle in the scanner. Images were selected to include two clusters each of animate and inanimate objects, which were counterbalanced into two clusters of real-world shape: those with a high-aspect ratio, or “bar-like” shape, and those with a low-aspect ratio, or “blob-like”, shape (Figure 1B). This resulted in 4 subordinate clusters of 8 images each: pets, insects, tools, and vegetables. Stimulus presentation and control for all experiments in the study was via PC computers running the Psychophysical Toolbox package ^19^, along with custom code, in Matlab (The Mathworks).

**Figure 1.**
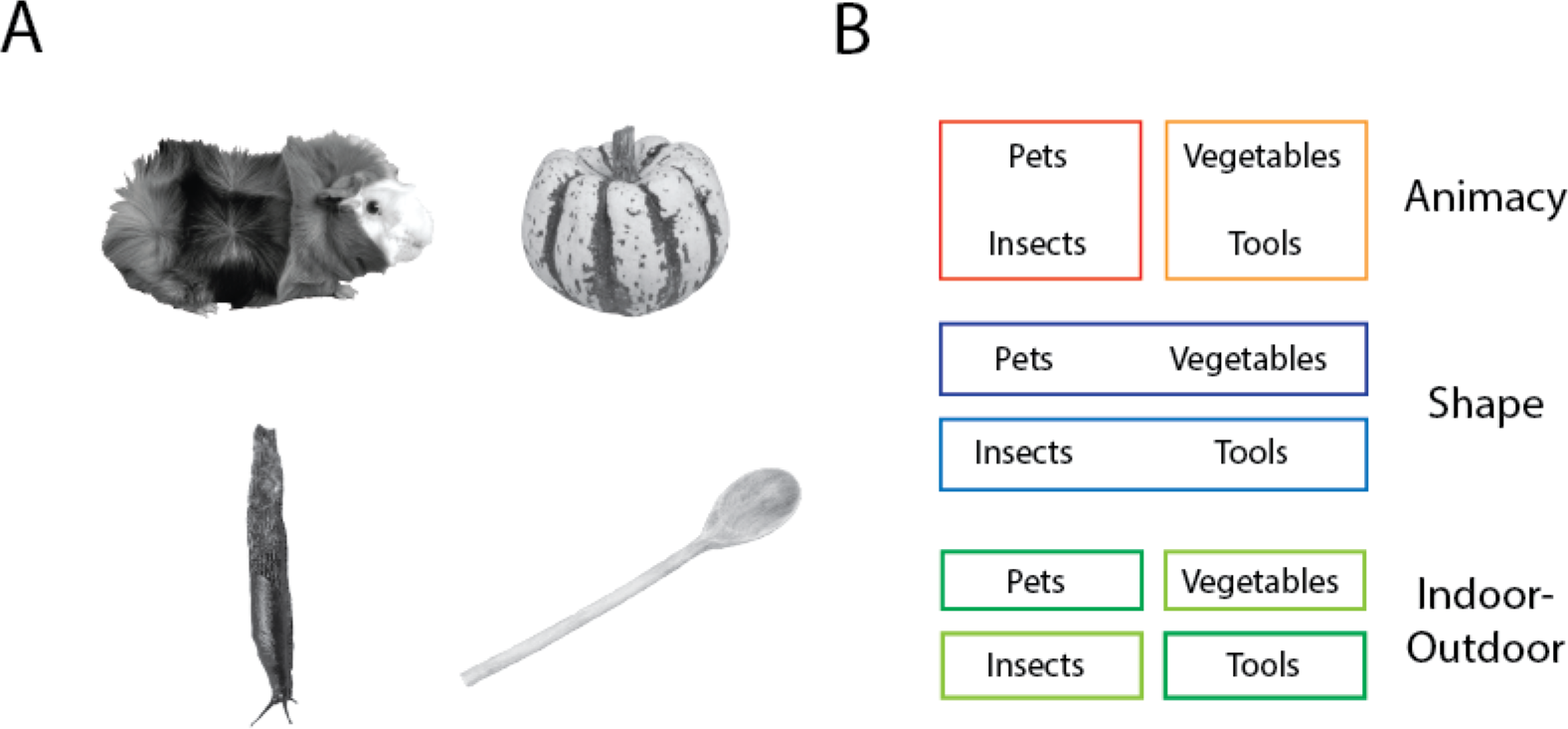
Stimuli and categorization task design. (A) Representative natural images of the pets, insects, vegetables, and tools stimuli. The sources for 3/4 of the displayed images are as follows: guinea pig (https://commons.wikimedia.org/wiki/File:AniarasKelpoKalle.jpg); slug (https://www.flickr.com/photos/brewbooks/2606728819); and wooden spoon (https://upload.wikimedia.org/wikipedia/commons/7/7b/Wooden_Spoon.jpg). These 3/4 images are published in compliance with a CC BY-SA license (https://creativecommons.org/licenses/by-sa/3.0/). (B) the category groupings of the stimuli for the animacy, shape, and indoor-outdoor tasks.

**Figure 2.**
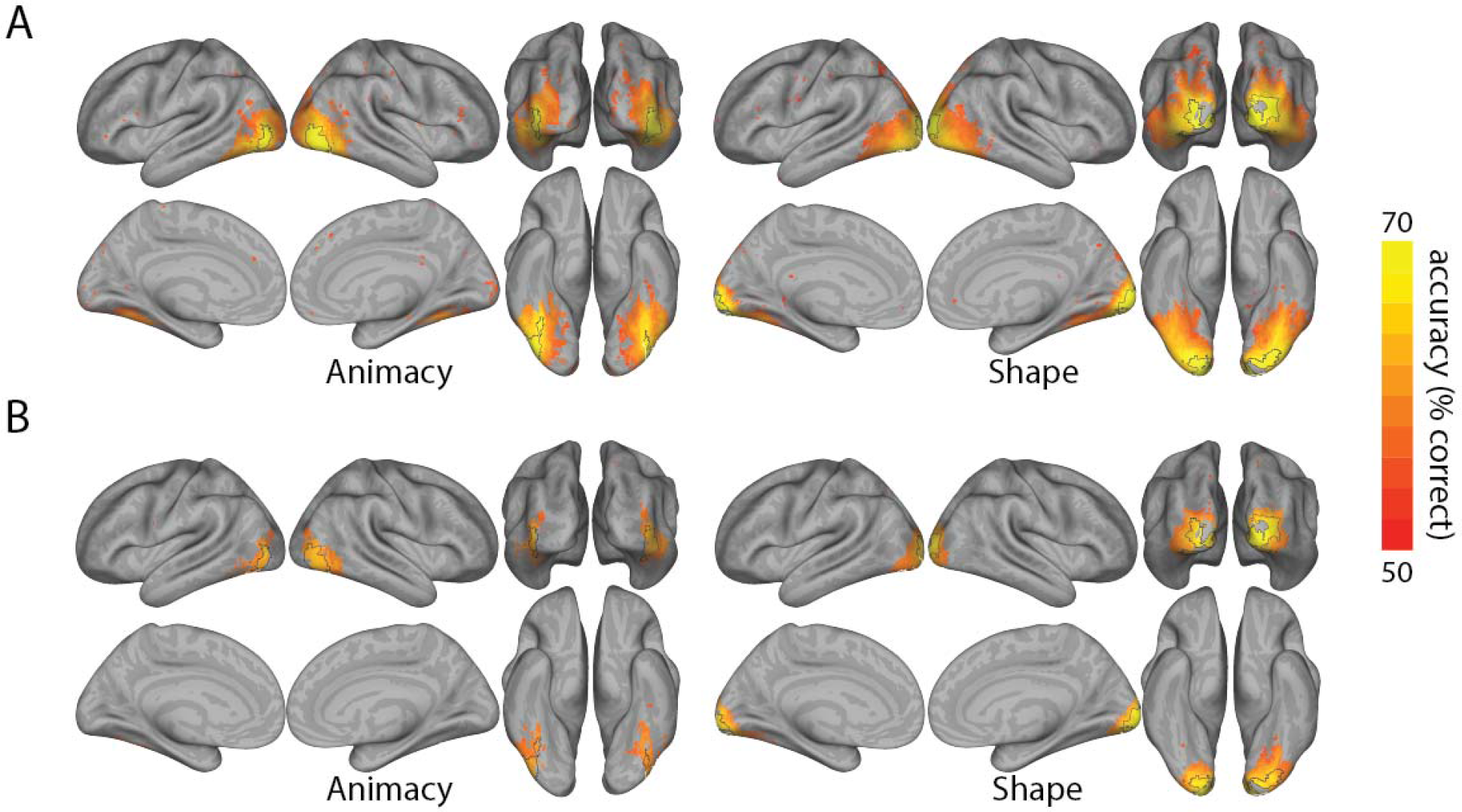
Decoding searchlight results. Results of searchlight analysis using standard leave-one-run-out cross-validation decoding (A), and cross-decoding (B), for both animacy and shape. Accuracy maps are projected onto the inflated cortical surface using CARET (van Essen et al. 2001), and thresholded at p < .005 (FDR-adjusted). Black lines on the surfaces indicate the location of the 10 % of voxels with highest classifier accuracy when using standard cross-validation (primarily in LOC for animacy, and in EVC for shape).

### Image analysis

To assess whether low-level image properties predicted stimulus animacy or shape, we applied the GIST model, which provides a summary of local orientation and spatial frequency information in an image, to our stimuli ^20^. Each image was segmented into a 4 × 4 grid, and Gabor filters with 8 different orientations and 4 spatial frequencies were applied independently to each grid block of the image. This procedure resulted in a 512 value descriptor vector (16 image blocks × 8 orientations × 4 spatial frequencies) for each image. We then applied a linear discriminant analysis (LDA) classifier to the descriptor vectors to carry out cross-decoding for the shape and animacy labels of the images. For example, a classifier trained to discriminate the animacy of (shape-matched) insects and tools images was tested on the (shape-matched) pets and vegetable images, and vice versa. This analysis allowed us to assess whether any information about animacy category or real-world shape generalized across stimulus clustering along the orthogonal dimension.

### Categorization tasks

Subjects performed three tasks in the scanner: an animacy task (is the object in the image “capable of self-movement”); a shape task (is it more “bar-like or blob-like”); and a contrived “indoor-outdoor” task, which crisscrossed the animacy and shape dimensions (Figure 1B). For the indoor-outdoor task it was emphasized to participants that one feature of some of the objects is that they tend to be found or kept indoors. Specifically, the tool stimuli were kitchen and bathroom items, and the pets (e.g. guinea pigs) tend to be kept in cages indoors. In contrast, we highlighted that the vegetables in our images are typically grown outdoors in the garden, and that the insects in our images are also found outside in trees or gardens. Thus, for the indoor-outdoor task, subjects were instructed to use a binary rule for categorizing the object images into groups that crisscrossed the animacy and shape divisions, but without overt reference to nonlinear decision rule. All three tasks were explained to subjects prior to scanning. Dependent measures for these tasks were observer choice responses and reaction times.

### Scanning procedure

Scanning consisted of a single sessions of 9 functional scans, followed by an anatomical scan. Using a rapid event-related design, each run consisted of a random sequence of trials including two repeats of each of the 32 stimuli and 11 fixation trials, for a total of 86 trials per run. Each stimulus trial began with the stimulus being presented for 1000 ms, followed by 2000 ms of fixation. The duration of each run was 4 m 40 s. During scanning, subjects first performed either the animacy then shape, or shape then animacy task, followed by the indoor-outdoor task, for 3 runs per task. Responses were made using the middle and index finger of the right hand. Since stimulus durations were longer than is typical when collecting RTs, participants were instructed to respond while the stimulus was still on the screen. To control for possible motor confounds, the response-button mapping was counterbalanced between runs.

### Imaging parameters

Data acquisition was carried out using a 3T Philips scanner, with 32-channel coil, at the Department of Radiology of the KU Leuven university hospitals. Functional MRI volumes were acquired using a 2D T2*-weighted echo planar (EPI) sequence: TR = 2 s; TE = 30 ms; FA = 90 deg; FoV = 216; voxel size = 3 × 3 × 3 mm; matrix size = 72 × 72. Each volume consisted of 37 axial slices aligned to encompass as much of the cortex as possible, with no gap. The T1-weighter anatomical volumes were acquired with an MP-RAGE sequence, 1 × 1 × 1 mm resolution.

### fMRI preprocessing and analysis

Preprocessing and analysis of the MRI data was carried out using SPM12 (v. 6906). For each participant, fMRI data was slice-time corrected, motion corrected, coregistered to the individual anatomical scan, normalized to standard MNI space, and then smoothed using a Gaussian kernel, 6 mm FWHM ^21^. After preprocessing the BOLD signal for each stimulus, at each voxel, was modeled using a GLM. The predictors for the model consisted of the 32 stimulus conditions and 6 motion correction parameters (translation and rotation along the *x*, *y*, and *z* axes). The time course for each predictor was characterized as a boxcar function convolved with the canonical hemodynamic response function. The GLM analysis produced one parameter estimate for each voxel, for each predictor, for each run.

### Decoding searchlight analysis

To determine where information about category and shape might be located, we carried out a volume-based decoding searchlight analysis (100 voxel neighborhoods) using LDA classifiers, as implemented in the CoSMoMVPA toolbox ^22^. The analysis was performed using two different cross-validation procedures. First, we utilized standard leave-one-run-out cross-validation. Second, mirroring the image analysis, we performed a cross-decoding searchlight analysis ^6^. To control for multiple comparisons, all p-values were FDR adjusted. To isolate neural features for carrying out NDBA, we selected the 10% highest classifying voxels within the area that showed significant decoding (see below).

### Neural distance-to-bound analysis

Following on previous work using NDBA we correlated observer RTs with neural distances from a classifier decision boundary through activation space. According to distance-to-bound models of discrimination and categorization, such as signal detection theory and its multidimensional extension ^12,13^, evidence quality increases with distance from a decision boundary through psychological space. Furthermore, RTs tend to decrease as the quality of evidence for a decision increases. Therefore, if differences in the latencies of RTs across stimuli are primarily the result of differences in the quality of the evidence they afford, then RTs for the stimuli should negatively correlate with the distance of stimulus representations from the decision boundary in the psychological space. NDBA applies the same logic to activation spaces in order to predict behavior ^14^. If an activation space carries information that is relied on by an observer performing a categorization task, then one can use a linear classifier applied to the space to estimate a decision boundary, and then correlate the distances of the neural responses of stimuli from this boundary with observer RTs for the same stimulus conditions.

Besides RTs, we have also found that neural distances negatively correlate with observer accuracies on animacy tasks, and that the drift rate parameter of an evidence accumulation model, which jointly models both dependent variables, can produce a larger correlation with neural distance ^18^. A simpler method for combining the strengths of both measures into a single value is via an efficiency score, which assumes a linear relationship between choice speed and accuracy. We opted to use the linearly integrated speed-accuracy score (LISAS), which respects this linear relationship and constitutes a form of accuracy (i.e. proportion error) adjusted RT ^23^. After computing this value for each stimulus for each subject, we group averaged the scores to generate a single behavioral vector to correlate with the individual subject neural distances. As a proxy for neural distances we used decision weights of an LDA classifier, averaged across cross-validation folds with a 4/5 train-test split. These values were calculated from regions identified by the 10% highest classifying voxels as determined by the searchlight analysis for animacy and shape, since RT-distance effect size largely tracks decodability across spatial and temporal features ^16,17^.

Since it has previously been observed that the RT-distance relationship for animacy tasks is primarily driven by animate stimuli we separately correlated the distances for each of the two categories for each of the three tasks ^17^. Crucially, for the indoor-outdoor task, we separately scaled the neural distances from the animacy and shape classifier boundaries for the high classifying voxels to percentiles, so that both sets of distances would receive equal weight. These distances were then summed, following the rationale of city block distance metrics assuming shape and animacy are independent dimensions. These scaled and summed values were then correlated with the adjusted RTs. Thus the neural distances for the indoor-outdoor task reflected information about both animacy and shape, which were crisscrossed dimensions in our stimulus design.

While in previous work we have used rank-order correlations in the present study we linearly correlated the neural distance and adjusted RT vectors, since this allowed us to adopt a joint measure of reliability from psychometrics that has been used in previous electrophysiological and neuroimaging studies in monkeys ^24,25^. For this we computed the mean split-half correlations for both adjusted RTs and neural distances, and then applied the Spearman-Brown formula to estimate the reliability of the full data sets. The joint reliability is then the square root of the product of the two coefficients ^26^. Since these values are positive, but the correlations between adjusted RTs and neural distances are predicted to be negative, the sign-value of the reliability estimates were inverted for visualization.

## Results

### Behavioral performance on categorization tasks

Participants performed three categorization tasks for the natural image stimuli while lying in an fMRI scanner. For the animacy task participants were highly accurate in their choice responses (mean accuracy = .91 ± .09), and were less accurate for the shape task (mean accuracy = .85 ± .11), and especially the indoor-outdoor task (mean accuracy = .81 ± .14). For RTs participants were fastest at the shape task (median RTs = 818 ms, ± 8 ms), than the animacy task (median RTs = 840 ms, ± 10 ms), and slowest for the indoor-outdoor task (median RTs = 896 ms, ± 10 ms). This pattern of behavioral results is consistent with the fact that subjects reported the indoor-outdoor task to be the most difficult, which was to be anticipated based on the contrived nature of the category division when applied to the stimulus images. Notably, the data of one participant, who was at chance for the indoor-outdoor task, was excluded from all further analysis.

In preparation for NDBA, accuracy and RT date were combined. Adjusting RTs using observer accuracy assumes a negative correlation between the two measures ^23^. This was generally observed across tasks, albeit only significantly for the animacy (12/15 participants, mean r = −.22 ± .25, t(14) = 03.33, p < .01) and shape tasks (10/15 participants, mean r = −.18 ± .29, t(14) = − 2.5, p < .05), but not the indoor-outdoor task (10/15 participants, mean r = −.11 ± .24, t(14) = −1.8, p = .09).

### Decoding searchlight for object animacy and shape

Decoding searchlight analysis was performed to find isolate high classifying voxels for object animacy and shape (Figure 1C). This revealed animacy-related information in the lateral occipital complex (LOC) and ventral temporal cortex (VTC), but crucially not early visual cortex (EVC). For shape information decodable information was found across the occipital lobe including EVC, as well as LOC and VTC. However, the highest 10 % classifying voxels for these analyses did not overlap, and were primarily in LOC for animacy, and EVC for shape. Image analysis showed that a cross-decoding classifier could readily distinguish between shape-types based on GIST descriptors (mean accuracy = .88), but crucially, not animacy (mean accuracy = .38), suggesting low-level properties of the images contained little category identifying information. In contrast, cross-decoding searchlight showed significant decoding for both animacy and shape, with significant clusters that tended to overlap with the regions of peak decoding using standard cross-validation (Figure 1D).

### Neural distance predicts categorization behavior across tasks

First, we correlated the adjusted RTs for the animacy task with neural distances from an animacy classifier boundary in LOC and EVC. In LOC there was a robust RT-distance correlation for the animate images (mean r = −.58, t(14) = −13.02, p < .001), but only a borderline significant effect for the inanimate images (mean r = −.12, t(14) = −2.08, p = .06). The difference between the mean correlations for the two categories was also highly significant (t(14) = −5.40, p < .001). Furthermore, the joint measures of reliability (grey bars in Fig 1E) suggest that there is explainable variance for the inanimate stimuli, but it is not well-captured by neural distance. In EVC there was a small but significant RT-distance correlation (mean r = −.13, t(14) = −2.58, p < .05) for the animate stimuli but not the inanimate stimuli (mean r = .08, t(14) = 1.10, p = .29), even though animacy could not be decoded from this region.

Second, we also calculated the correlations between adjusted RT neural distance for our shape task. There were significant negative correlations for both the blob-like (mean r = −.32, t(14) = − 4.50, p < .01) and bar-like stimuli (mean r = −.17, t(14) = −3.03, p < .01) in EVC, where there was greatest shape decoding, with no significant difference between the effects. A significant negative correlation was also observed in LOC for the bar-like stimuli (mean r = −.16, t(14) = −3.34, p < .01), but not for the blob-like category (mean r = −.10, t(14) = −1.46, p = .17), even though the joint reliability for the blob-like images was similar for both ROIs. In contrast, for the bar-like stimuli the correlation in LOC was close to the noise ceiling, which was much lower than in EVC.

Third, we carried out the same analysis for the crisscrossing indoor-outdoor task. Here we observed no effects in EVC for either the indoor (mean r = .10, t(14) = 1.84, p = .09) or outdoor (mean r = −.02, t(14) = −.21, p = .84) category stimuli. There was an effect for the indoor stimuli in LOC (mean r = −.12, t(14) = −3.04, p < .01) but not the outdoor stimuli (mean r = .00, t(14) = −.01, p = .99). With the exception of the one significant effect, the individual data points were highly variable with the mean correlation coefficients even trending in the wrong, positive direction. Overall, the joint reliability was also lower for the indoor-outdoor task than for the other two tasks. Furthermore, while

## Discussion

In the present study we sought to determine whether neural distance would predict observer RTs when stimuli are balanced along orthogonal object category and shape dimensions. To this end, we correlated neural distances from classifier decision boundaries for object animacy and shape in EVC and LOC with accuracy adjusted RTs from three categorization tasks: animacy, shape, and indoor-outdoor. In each case we observed, for at least one side of the category divisions, the predicted negative correlation between neural distance and adjusted RTs.

Stimulus animacy was highly decodable from LOC, consistent with previous findings ^27–29^. Also, as cross-decoding was used to both investigate whether category information can be discriminated by image properties and to isolate category-specific information in the ventral pathway ^5,6,30^, our results cannot be explained by the sorts of image properties captured by GIST ^2–4^. When neural distances from this region were correlated with adjusted RTs we found a strong RT-distance effect for the animate, but not inanimate, exemplars. This same pattern of results has been observed in previous studies using NDBA to investigate the animacy division. In particular, Carlson *et al.* ^15^ found a robust RT-distance effect in human VTC, which was driven entirely by the animate exemplars, and taking a searchlight approach to NDBA Grootswagers *et al.*^17^ found that neural distance for animate exemplars correlated with RTs in LOC and VTC, and largely tracked the areas of peak decoding. The same asymmetry has also been observed when applying NDBA to human MEG time-series data ^16,18^. The fact that a weak RT-distance effect was also observed for animate images in ECV is also consistent with previous findings of Carlson *et al.*^15^, and suggests that the visual properties represented in the region influence how easily a stimulus can be categorized even in situations in which these low-level properties by themselves are not sufficient for categorization.

The present study differed from previous studies using NDBA in three important respects. First, unlike in these previous studies, we controlled shape properties so that overall the animacy RT-distance effects cannot be easily explained by low-level properties ^8^. For this reason, our results provide more compelling evidence that observers are recruiting category-related information encoded in high-level visual cortex when performing the animacy categorization task.

Second, previously we have found that observer choice accuracies also correlate with neural distance. When both accuracy and RT distributions are jointly modeled using evidence accumulation models, this provides a more complete description of the observer performance, and the drift rate parameters of these models can also be correlated with neural distance ^18^. Efficiency scores provide a simpler method for combining both measures, and in the present study we used the recently proposed LISAS ^23^. Future studies might systematically compare whether drift rates of evidence accumulation models, which require more intensive procedures for parameter estimation, in fact show superior effects to the analytically simple efficiency scores when carrying out NDBA.

Third, an important consideration in analyses that aim to characterize the structure of activation spaces of brain regions is to have an estimate of data reliability. This is common practice when carrying out representational similarity analysis, or RSA ^31,32^. In particular, cross-validated distances may provide the best technique for estimating the neural dissimilarity relationships at the heart of RSA, and data reliability can be used to estimate a noise ceiling for correlations between model and neural dissimilarity matrices. To date, previous studies using NDBA have not used the same quality controls in estimating neural distances from a decision boundary. In the present study we rectified this by cross-validating our estimates of neural distance and using a joint measure of both behavioral and neural reliability to provide un upper bound on effect sizes ^24,25^. Notably, the same joint measure of reliability could be employed in RSA studies where linear correlation coefficients can be used to compare dissimilarity matrices of neural and behavioral data. The importance of including such reliability estimates in the current study is illustrated by the large variation in reliability in Figure 3; even between analyses that target the same brain area. This shows, for example, that the relatively low correlation between RTs and neural distance for elongated shapes can mostly be predicted from a low reliability of the data that are correlated. In contrast, the low correlation for inanimate objects is much lower than what would be predicted based upon the reliability of the data.

**Figure 3.**
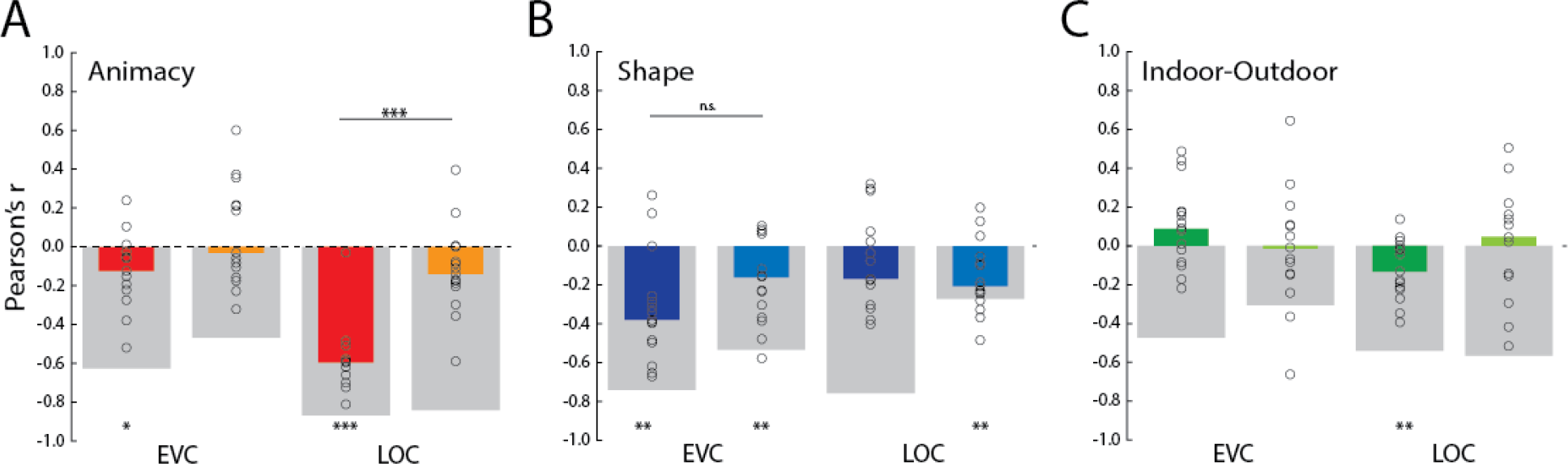
Neural distance-to-bound results. Results of neural distance-to-bound analysis for each of the three categorization tasks: animacy (A); shape (B): and indoor-outdoor (C). Color coding of median bars reflects the stimulus groupings for each task from Figure 1B. Neural distances are for EVC and LOC features, based on the highest classifying voxels from the searchlight analysis (Figure 1C). Thick gray bars are the inverted joint reliability of the neural distances and accuracy adjusted RTs. * = p, .05; ** = p < .01; *** = p < .001.

It is also useful to compare our results to recent studies using stimulus designs that orthogonalized object category and shape and observed information about both stimulus properties in the same brain regions. First, Kaiser *et al.*^6^ used cross-decoding searchlight to locate information about shape matched images of clothing (gloves and shirts) and body parts (hands and torsos). Cross-decoding revealed information about shape in EVC but not category, with overlapping clusters for both in LOC. Second, Bracci and Op de Beeck^7^ used a more complex design of 6 object categories and 9 shape types, and found information for both stimulus properties was present in multiple sub-regions of LOC and VTC. In contrast to these studies, we only observed an overlap in object category and shape information in LOC and VTC when carrying out a searchlight analysis with standard cross-validation. When using cross-decoding, information about shape was confined to EVC, while information about animacy was still found in LOC and to a lesser extent VTC; though a small RT-distance effect was still observed in LOC for the bar-like objects. In our stimulus design high and low-level shape are less conflated for the bar-like stimuli, which might account for why there is still a shape effect in LOC. More generally, an important difference between the present study and these previous findings is that low- and high-level shape were not differentiated in their stimulus designs.

The results for the indoor-outdoor task are difficult to interpret, but suggests that representations related to object category and shape might first be recruited, and the timing in the activation of these representations then impact performance on the indoor-outdoor task. It is also notable that both clusters for the indoor category, pets and tools, showed significant effects in LOC. This may be because pets may produce a more decodable animacy response than insects ^33^, and there is evidence that sub-regions of LOC code specifically for tools ^34,35^. Importantly, the size of the significant correlations in the indoor-outdoor task is much smaller than some of the correlations in the two other tasks, in absolute terms as well as relative to the reliability (explainable variance).

In summary, previous studies have used neural distance to predict observer behavior on visual categorization tasks for animacy. We built on these findings using a stimulus design that orthogonalized object category and shape ^8^. In line with previous results, we observed a robust effect for animacy in LOC driven by the animate stimuli, as well as a shape effect in EVC, and a smaller crisscross effect using a more contrived category structure that crisscrossed the two dimensions. Taken together these results suggest there is potential to expand the neural distance-to-bound approach to other divisions beyond animacy and object category per se ^17^.

## Author contribution statement

Both authors jointly designed the experiment. J.B.R collected and analyzed the data, and wrote the main manuscript and prepared the figures. Both authors reviewed the manuscript.

## Acknowledgements

This work was supported by the European Research Council (ERC-2011-StG-284101), a federal research action (IUAP-P7/11), a Hercules grant ZW11_10, and a KU Leuven research council grant (C14/16/031) to H.O. This project has received funding from the FWO and European Union’s Horizon 2020 research and innovation programme under the Marie Skłodowska-Curie grant agreement No 665501, via a FWO [PEGASUS]^2^ Marie Sklodowska-Curie fellowship (12T9217N) to J.B.R.

